# Regulation of inositol 5-phosphatase activity by the C2 domain of SHIP1 and SHIP2

**DOI:** 10.1101/2023.07.31.551266

**Authors:** William J Bradshaw, Emma C Kennedy, Tiago Moreira, Luke A Smith, Rod Chalk, Vittorio L Katis, Justin LP Benesch, Paul E Brennan, Emma J Murphy, Opher Gileadi

## Abstract

SHIP1, an inositol 5-phosphatase, plays a central role in cellular signalling. As such, it has been implicated in many conditions. Exploiting SHIP1 as a drug target will require structural knowledge and the design of selective small molecules. We have determined apo, and magnesium and phosphate-bound structures of the phosphatase and C2 domains of SHIP1. The C2 domains of SHIP1 and the related SHIP2 modulate the activity of the phosphatase domain. To understand the mechanism, we performed activity assays, hydrogen-deuterium-exchange mass spectrometry and molecular dynamics on SHIP1 and SHIP2. Our findings demonstrate that the influence of the C2 domain is more pronounced for SHIP2 than SHIP1. We determined 91 structures of SHIP1 with fragments bound, with some near the interface between the two domains. We performed a mass spectrometry screen and determined four structures with covalent fragments. These structures could act as starting points for the development of potent, selective probes.

## Introduction

SH2-containing-inositol-5-phosphatases (SHIP1 and SHIP2) are key parts of the PI3K/AKT/mTOR signalling pathway. SHIPs dephosphorylate phosphatidylinositol-3,4,5-trisphosphate (PI(3,4,5)P_3_) to produce phosphatidylinositol-3,4-bisphosphate (PI(3,4)P_2_). The balance between these two second messengers and related inositol- and phosphatidylinositol phosphates (PIPs) ultimately affects a wide number of cellular processes including inflammation, autophagy, cell migration, metabolism and cell growth.^1, 2^ Therefore, it is not surprising that SHIPs have been implicated in many conditions, including a range of cancers,^3–5^ diabetes,^6^ hypertension,^7^ and graft versus host disease.^8^

Genome wide association studies and follow-up investigations have demonstrated a link between SNPs rs10933431 in intron 2 and rs35349669 in intron 10 of the *INPP5D* gene (SHIP1), increased expression of SHIP1, and increased risk of late-onset Alzheimer’s disease (LOAD) in Caucasians.^9–11^ The latter of these two mutations, on the other hand, does not appear to result in increased risk of LOAD in the Han Chinese population.^12^

The link between SHIP1 and Alzheimer’s disease is believed to arise through its role in inflammatory processes in the brain, specifically through the regulation of microglia and cytokine release via signalling from TREM2 and DAP12.^13–17^ The level of expression of the two SHIPs varies between tissues.^18, 19^ In the brain specifically, *INPP5D* expression in enriched in microglia, with lower levels in other cells. SHIP2 (*INPPL1*) has a lower level of expression but this is more consistent across a wider range of cells, including microglia and astrocytes.^20^ Some doubt has recently been cast on whether the link to Alzheimer’s disease through microglial function is unique to SHIP1, or involves both SHIPs.^21^ Further complicating matters, SHIPs have roles that extend beyond their phosphatase activity, as they possess ubiquitin and SH3 binding sites, and their phosphatase-dependent and -independent roles overlap.^22, 23^

Clearly, an improved understanding of how the two SHIPs carry out their functions and their respective roles in Alzheimer’s disease and other conditions is required. To this end, a number of inhibitors and activators that can be used as probes have been developed in recent years, some of which are reported to have near-equal efficacy on the two SHIPs, while others have been reported as having up to 20-fold selectivity.^21, 24, 25^ The utility of many of these compounds is somewhat limited and better chemical probes, particularly ones that have been structurally engineered to have high affinities and to be selective, would prove invaluable.

The SHIPs possess an N-terminal SH2 domain, which mediates interactions with phosphorylated membrane proteins such as c-kit, DAP12 and several others.^14, 26, 27^ This is followed by a disordered region of approximately 60 residues and a Dimerisation/RhoA binding domain.^28, 29^ Centrally, they have a plekstrin homology (PH) domain followed by a 5-phosphatase domain, a C2 domain and a disordered region of approximately 300 residues containing the aforementioned SH3 binding sites. SHIP2 has a C-terminal ubiquitin interacting motif and a SAM domain.^23^ The product of SHIP catalysis, PI(3,4)P_2_, is believed to act as an allosteric activator of the inositol phosphatase activity, the C2 domain has been demonstrated to be essential to this allosteric effect: activators only work when the C2 domain is present.^30^

The human genome codes for 10 inositol 5-phosphatases. Structures have so far been determined for 5 of these: INPP5B, OCRL, SHIP2,^31^ synaptojanin-1,^32^ and INPP5E (2XSW, Trésaugues et al., unpublished). The active site is well conserved between these proteins but there are a small number of differences that could be exploited if selective inhibitors of inositol 5-phosphatases are to be developed. The structure of the phosphatase domain of SHIP2 with the C2 domain has also been determined.^33^ The authors notably highlighted four loops (loop 1 to loop 4) in the phosphatase domain of SHIP2, and suggested roles for three of them. Loop 2 (K510 to G518 in SHIP1, K531 to G539 in SHIP2) was proposed to penetrate the membrane and deliver the substrate to the active site. Loop 3 (D566 to N573 in SHIP1, D587 to D594 in SHIP2) interacts with the C2 domain and is thought to be important to transmitting the allosteric effects of the C2 domain to the active site in the phosphatase domain. Loop 4 (T656 to N666 in SHIP1, N674 to N684 in SHIP2) forms the final part of this allosteric conveyor and also forms part of the active site. Le Coq *et al*. proposed a mechanism of communication between the C2 and phosphatase domains through structures, activity assays on several mutants, and molecular dynamics.

Here, we present two high-resolution structures of the phosphatase and C2 domains of SHIP1: one in its apo form and another with magnesium and phosphate bound to the active site. Comparison to the structures of other inositol 5-phosphatases reveals differences that are likely to be crucial for the development of novel and selective compounds. We have explored the role of the C2 domain and the allosteric mechanism through activity assays, hydrogen-deuterium exchange mass spectrometry experiments (HDX-MS) and molecular dynamics simulations. These results provide evidence that the C2 domain is particularly important for SHIP2 activity, but less so for SHIP1. We also present the results of a SHIP1 crystallography-based fragment screen, with 91 compounds bound that may serve as starting points for the development of novel inhibitors or degraders. As a follow-up to this, we performed a mass spectrometry (MS) screen of covalent binders and have determined a further 4 high-resolution structures with covalent compounds bound.

## Results

### The structure of SHIP1

We have determined the apo-structure of the phosphatase and C2 domains of human SHIP1 to a resolution of 1.48 Å and refined it to R-factors of 0.173 and 0.201. A magnesium and phosphate-bound structure of the same domains has also been determined with an elliptical high-resolution cut-off selected at resolutions between 1.34 Å and 1.09 Å. This was refined to R-factors of 0.136 and 0.157. Each structure has one chain in the asymmetric unit and the two structures superpose on each other with an RMSD of 0.17 Å (Figure 1). The structure is very similar to that of mouse SHIP1, which was determined at approximately the same time,^34^ to which our two structures have RMSDs of 0.29 Å (apo) and 0.27 Å (MgPO_4_) and an identity of 95%. The fold is also very similar to that of the phosphatase and C2 domains of human SHIP2 with the two structures superposing on the 8 chains of 5OKM^33^ with RMSDs between 0.59 Å and 0.80 Å, with an identity of 57%. The magnesium and phosphate-bound structure is very similar to 5OKN, the equivalent SHIP2 structure, which has a D607A active site mutation.^33^ However, the significantly higher resolution of the present structure (1.34-1.09 Å vs 2.65 Å), allows more precise positioning of the magnesium and phosphate ions, and orientation the latter, within the active site and all waters bound to them, clarifying precise interactions within the active site and their importance in catalysis (Figure 2). Most differences between the two structures can be put down to limitations of the resolution of the SHIP2 structure. There are however, some important differences between our magnesium and phosphate bound structure, the magnesium and phosphate bound OCRL structure (4CMN)^31^ and the substrate bound Synaptojanin-1 structure.^32^ All structures presented here assume a loop 4-in conformation. This can be compared to structures of SHIP2 exhibiting either loop 4-in or loop 4-out conformations (Figure 2E), both of which are believed to be important to the catalytic mechanism and regulation of phosphatase activity by the C2 domain. While the conformation of this loop does differ somewhat from that seen in the SHIP2 loop 4-in structure, 3NR8^31^, care should be taken in over interpreting any differences due to potential packing artefacts in the present structures and in the SHIP2 structure.

**Figure 1.**
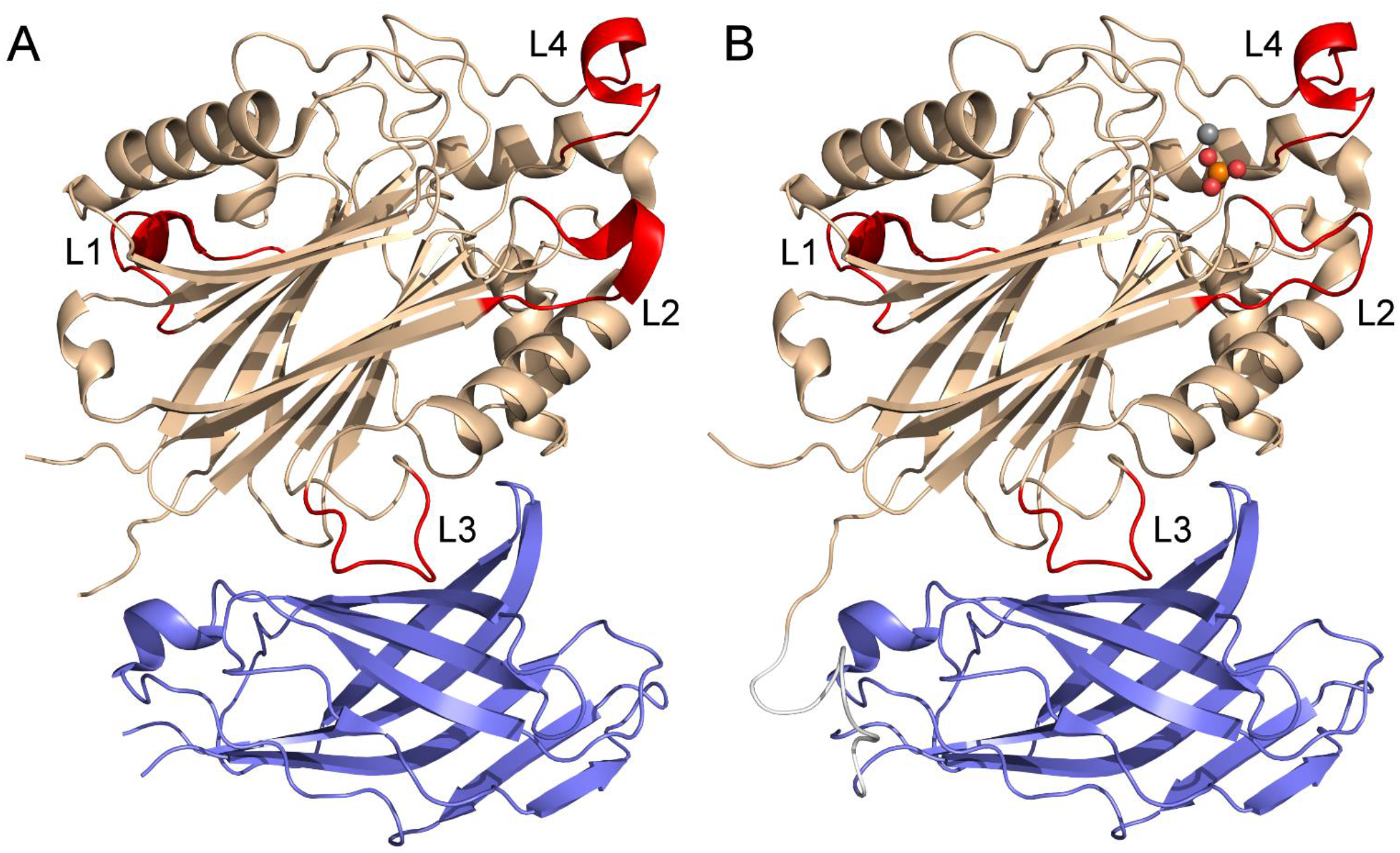
The structure of the phosphatase and C2 domains of SHIP1. **(A)** The apo structure. The phosphatase domain is coloured brown and the C2 domain is blue. Loops 1 to 4 of the phosphatase domain, as numbered in Le Coq et al. (2017) for SHIP2, are coloured red. **(B)** The magnesium and phosphate bound structure. Domains are coloured as in **A**. A magnesium ion (grey) and a phosphate ion (orange and red) are shown bound to the active site.

**Figure 2.**
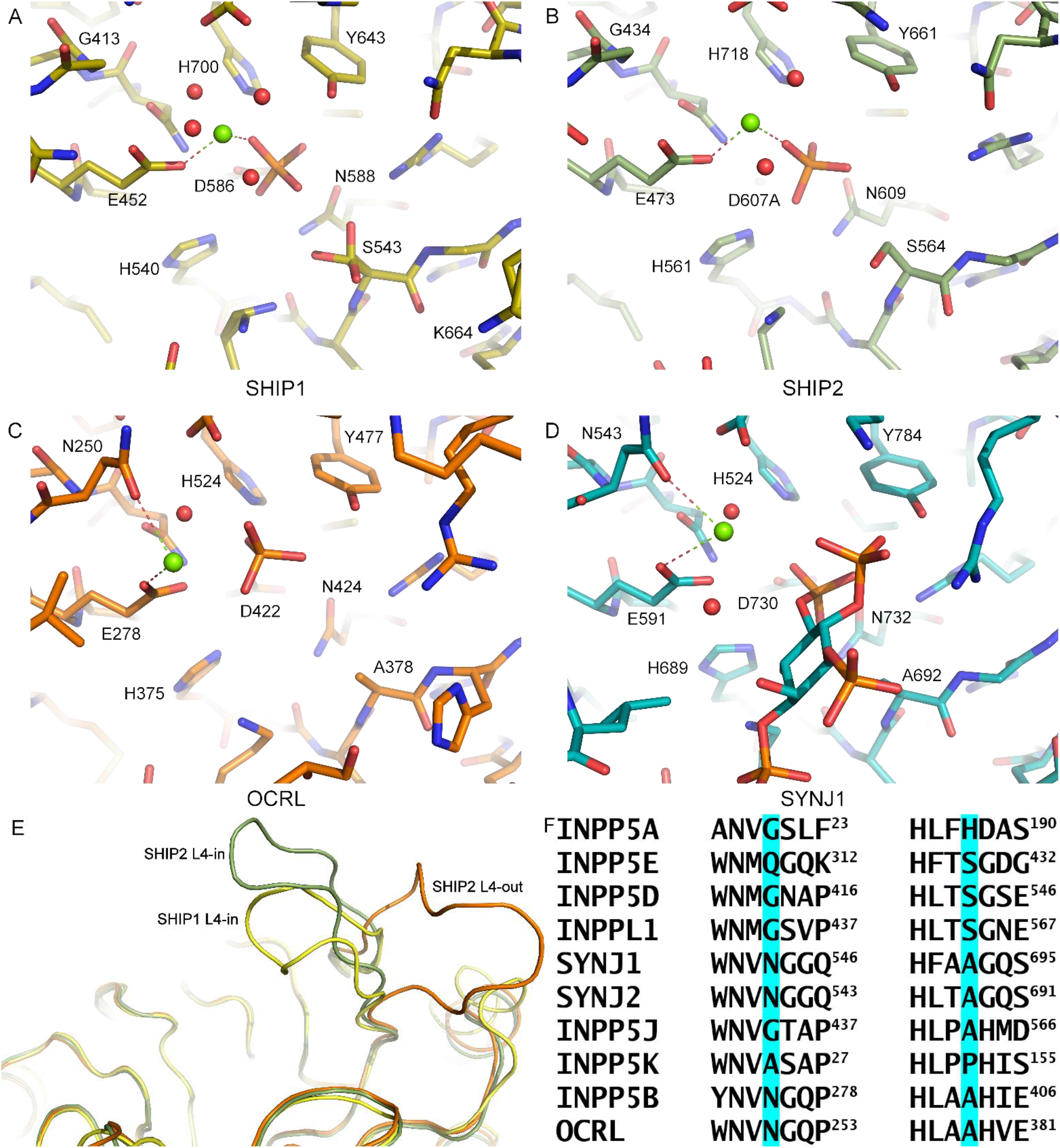
Active site comparison between INPP5 family members. **(A)** Magnesium and phosphate bound to the present SHIP1 structure (6XY7). The magnesium is octahedrally coordinated by E542, the phosphate and 6 waters. The phosphate is also bound to H540, S543, D586, N588 and H700. **(B)** Magnesium and phosphate bound to SHIP2 D607A (6OKN).^33^ **(C)** Magnesium and phosphate bound to OCRL (4CMN).^31^ As well as being coordinated by the phosphate and E278 (equivalent to E452 in SHIP1), the magnesium ion is also coordinated by N250, which is replaced by a glycine in the SHIPs. The phosphate is now bound to E278, Y477 and H524 (equivalent to E542, Y643 and H700 in SHIP1). **(D)** Magnesium and PI(3,4,5)P_3_-DiC8 bound to Synaptojanin 1 (7A17).^32^ Magnesium binding resembles that of OCRL, while phosphate binding resembles that of the SHIPs, but with space for an attacking water. **(E)** Comparison of loop 4 conformations between SHIP1 and SHIP2. Loop 4 is shown above the active site. The SHIP1 L4-in conformation shown is from the magnesium and phosphate bound structure and is yellow. The SHIP2 L4-in conformation is 3NR8 chain B, shown in green.^31^ The SHIP2 L4-out conformation is 5OKM chain B, shown in orange.^33^ (F) Alignment of portions of the active site in each INPP5. Gly413 and Ser543 in SHIP1 and their equivalents in other INPP5s are highlighted in blue.

### Activity assays

To determine the catalytic activity of SHIP1 and to compare it to that of SHIP2, assessing the effect of the C2 domain on the activity of the phosphatase domain, we performed activity assays on constructs of both SHIP1 and SHIP2 coding for the phosphatase domain alone (Ptase) and the phosphatase and C2 domains (Ptase-C2) (Figure 3). An assay which detects the production of inorganic phosphate was developed using Phosphate Sensor, a fluorescently (MDCC) tagged *E. coli*-derived inorganic phosphate binding protein. This sensor is more sensitive than the commonly used malachite green with the lower limit of quantification in the picomolar range compared with the nanomolar range for malachite green, this allows accurate determination of initial rates at lower substrate concentrations. However, the increased sensitivity means phosphate contamination of the substrate limits the range of concentrations that can be used, therefore K_m_ can be difficult to measure accurately if affinity for the substrate is low. For each construct, K_m_ and k_cat_ were determined for PI(3,4,5)P_3_-DiC8 and I(1,3,4,5)P_4_ using the phosphate sensor assay (Table 2). For the Ptase-C2 constructs, the two proteins showed similar levels of activity for respective substrates, although for PI(3,4,5)P_3_-DiC8, the k_cat_ of SHIP1 was approximately 50% higher than SHIP2’s, while SHIP2 showed slightly higher activity for I(1,3,4,5)P_4_ than SHIP1 did. For both substrates, SHIP2 showed a large increase in k_cat_ when the C2 domain was present. Much more modest differences were observed in k_cat_ for SHIP1. For both substrates, SHIP1 showed a slight increase in K_m_ in the presence of the C2 domain. SHIP2 showed a decrease in K_m_, and therefore an increase in affinity for IP_4_ with the C2 domain. The error in the calculated K_m_ of SHIP2’s phosphatase domain alone for PIP_3_ was large due to very low activity, so no conclusion can be drawn about differences in K_m_.

**Figure 3.**
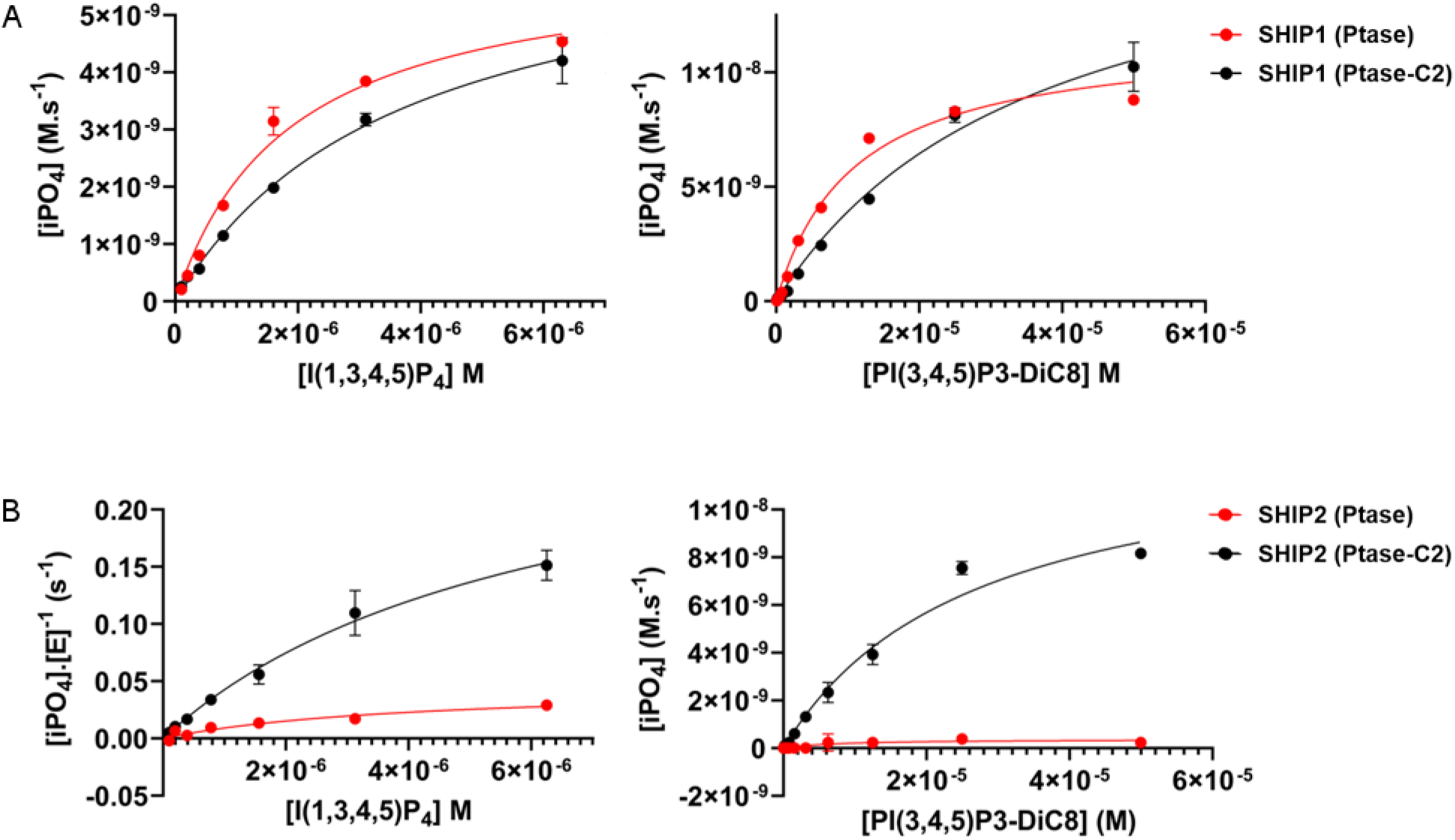
Activity assays. **(A)** Phosphatase activity of 50 nM SHIP1 against either I(1,3,4,5)P_4_ (left) or PI(3,4,5)P_3_-DiC8 (right). There are no significant differences in the catalytic activity in the presence (black circles) or absence of the C2 domain (red circles) for either substrate. **(B)** Phosphatase activity of 50 nM SHIP2 against either I(1,3,4,5)P_4_ (left) or PI(3,4,5)P_3_-DiC8 (right), as seen previously^33^ the catalytic efficiency of SHIP2 is greater in the presence of the C2 domain (black circles) than the phosphatase alone (red circles) (N.B. For I(1,3,4,5)P_4_ the SHIP2 (Ptase-C2) was at 15 nM and initial rates have been plotted as phosphate produced per second per molar concentration of enzyme). Data are represented as mean ± SEM.

**Table 1.** Crystallographic and refinement statistics outer shell statistics are given in brackets.

|  | Apo | Mg PO4 bound | Z2738285202 | Z1742148362 | Z1763271112 | Z56948267 |
| --- | --- | --- | --- | --- | --- | --- |
| Crystallographic statistics |  |  |  |  |  |  |
| Space group | P2<sub>1</sub>2<sub>1</sub>2<sub>1</sub> | P2<sub>1</sub>2<sub>1</sub>2<sub>1</sub> | P2<sub>1</sub>2<sub>1</sub>2<sub>1</sub> | P2<sub>1</sub>2<sub>1</sub>2<sub>1</sub> | P2<sub>1</sub>2<sub>1</sub>2<sub>1</sub> | P2<sub>1</sub>2<sub>1</sub>2<sub>1</sub> |
| Cell dimensions (Å, °) | 62.9 80.1 90.3<br>90 90 90 | 62.5 79.1 89.3<br>90 90 90 | 62.7 79.4 89.6<br>90 90 90 | 62.5 79.2 89.4<br>90 90 90 | 62.5 79.0 89.3<br>90 90 90 | 62.7 79.6 89.6<br>90 90 90 |
| Resolution (Å) | 80.1-1.48<br>(1.51-1.48) | 59.2-1.09<br>(1.14-1.09) | 79.4-1.40<br>(1.42-1.40) | 79.2-1.45<br>(1.47-1.45) | 62.5-1.30<br>(1.32-1.30) | 51.4-1.40<br>(1.42-1.40) |
| R<sub>merge</sub> | 0.187 (3.948) | 0.152 (2.190) | 0.170 (2.324) | 0.194 (3.638) | 0.096 (2.308) | 0.134 (2.517) |
| R<sub>pim</sub> | 0.062 (1.639) | 0.030 (0.480) | 0.070 (1.010) | 0.081 (1.495) | 0.040 (0.998) | 0.057 (1.095) |
| CC<sub>1/2</sub> | 0.997 (0.385) | 0.998 (0.601) | 0.998 (0.444) | 0.998 (0.453) | 1.000 (0.581) | 0.998 (0.422) |
| Mean <I/σI> | 11.4 (1.2) | 11.7 (1.4) | 8.5 (1.1) | 8.0 (0.7) | 11.6 (0.7) | 9.6 (1.1) |
| Completeness (%) |  |  |  |  |  |  |
| Spherical | 100.0 (99.3) | 87.6 (37.0) | 100.0 (100.0) | 100.0 (100.0) | 100.0 (100.0) | 100.0 (100.0) |
| Ellipsoidal | N/A | 94.2 (59.6) | N/A | N/A | N/A | N/A |
| Total reflections | 1,488,660<br>(49,808) | 4,301,505<br>(163,879) | 1,166,735<br>(54,505) | 1,021,758<br>(51,037) | 1,390,256<br>(65,597) | 1,064,015<br>(52,281) |
| Total unique reflections | 76,674 (3,736) | 159,826 (7,907) | 88,594 (4,478) | 79,287 (3,864) | 109,198 (5,345) | 88,912 (4,359) |
| Multiplicity | 19.4 (13.3) | 26.9 (20.7) | 13.2 (12.2) | 12.9 (13.2) | 12.7 (12.3) | 12.0 (12.0) |
| Refinement statistics |  |  |  |  |  |  |
| R<sub>work</sub>/R<sub>free</sub> | 0.173/0.201 | 0.136/0.157 | 0.137/0.190 | 0.144/0.218 | 0.144/0.191 | 0.133/0.190 |
| RMSDs |  |  |  |  |  |  |
| Bond lengths (Å) | 0.013 | 0.011 | 0.013 | 0.013 | 0.012 | 0.013 |
| Bond angles (°) | 1.791 | 1.702 | 1.785 | 1.753 | 1.718 | 1.770 |
| Ramachandran statistics (%) |  |  |  |  |  |  |
| Favoured | 97.3 | 97.8 | 97.6 | 95.0 | 96.3 | 94.4 |
| Allowed | 2.7 | 2.2 | 2.4 | 5.0 | 3.7 | 5.6 |
| Outliers | 0 | 0 | 0 | 0 | 0 | 0 |
| Average B-factors |  |  |  |  |  |  |
| Protein | 21.0 | 14.3 | 19.9 | 21.3 | 21.5 | 22.0 |
| Ligand | N/A | 7.7 | 22.9 | 26.8 | 31.1 | 26.9 |
| Water | 28.4 | 25.4 | 31.1 | 32.2 | 33.5 | 32.3 |
| Number of Atoms |  |  |  |  |  |  |
| Protein | 3742 | 4153 | 3871 | 3835 | 3936 | 3888 |
| Ligand | N/A | 6 | 16 | 13 | 13 | 9 |
| Water | 374 | 491 | 528 | 482 | 536 | 495 |
| PDB Code | 6IBD | 6XY7 | 8PDG | 8PDH | 8PDI | 8PDJ |

**Table 2.** Activity Assays. Errors are SEM

| Phosphate Sensor | PI(3,4,5)P<sub>3</sub>-DiC8 |  | I(1,3,4,5)P<sub>4</sub> |  |
| --- | --- | --- | --- | --- |
|  | K<sub>m</sub> (μM) | k<sub>cat</sub> (s<sup>-1</sup>) | K<sub>m</sub> (μM) | k<sub>cat</sub> (s<sup>-1</sup>) |
| SHIP1 Ptase | 11.4 ± 1.6 | 0.24 ± 0.01 | 1.8 ± 0.2 | 0.12 ± 0.006 |
| SHIP1 Ptase-C2 | 39.4 ± 5.7 | 0.38 ± 0.03 | 3.7 ± 0.4 | 0.13 ± 0.008 |
| SHIP2 Ptase | 19.3 ± 13* | 0.009 ± 0.003 | 6.90 ± 0.75 | 0.064 ± 0.003 |
| SHIP2 Ptase-C2 | 26.6 ± 3.7 | 0.26 ± 0.02 | 2.7 ± 0.6 | 0.18 ± 0.02 |

\*Very low activity, poor fit to Michaelis-Menten.

### Hydrogen-deuterium exchange

To characterise the effect of the C2 domain on flexibility of the phosphatase domain, hydrogen-deuterium exchange mass spectrometry was performed on the constructs of SHIP1 and SHIP2 with and without the C2 domain (Figure 4). For SHIP1 after digestion with pepsin, 88 peptides were detected that were common between the two constructs, covering 88% of the phosphatase domain. For SHIP2, 70 shared peptides were detected covering 86% of the phosphatase domain (Figure S1). Almost all differences in deuterium uptake (DU) that were observed between constructs with and without the C2 domain showed greater DU in the shorter construct, with differences being more prominent for SHIP2 than for SHIP1. That is to say, the phosphatase domains alone were more flexible/solvent accessible than the phosphatase domains with their C2 domains. This is to be expected in regions normally buried in the interface between the two domains, but potentially not in more remote parts of the phosphatase domain. The greatest disparities were observed in residues approximately between 580 and 650 in SHIP2, encompassing part of helix 4, loop 3 (which interacts with the C2 domain), strand 9 (connecting loop 3 to the active site) and helices 5, 6 and 7. Loop 3 and helix 7 in SHIP1 also showed the greatest degree of deprotection, but the effect on residues between these regions was less pronounced. Examples of regions where a statistically significant difference in DU was observed are shown in Figures S2 and S3.

**Figure 4.**
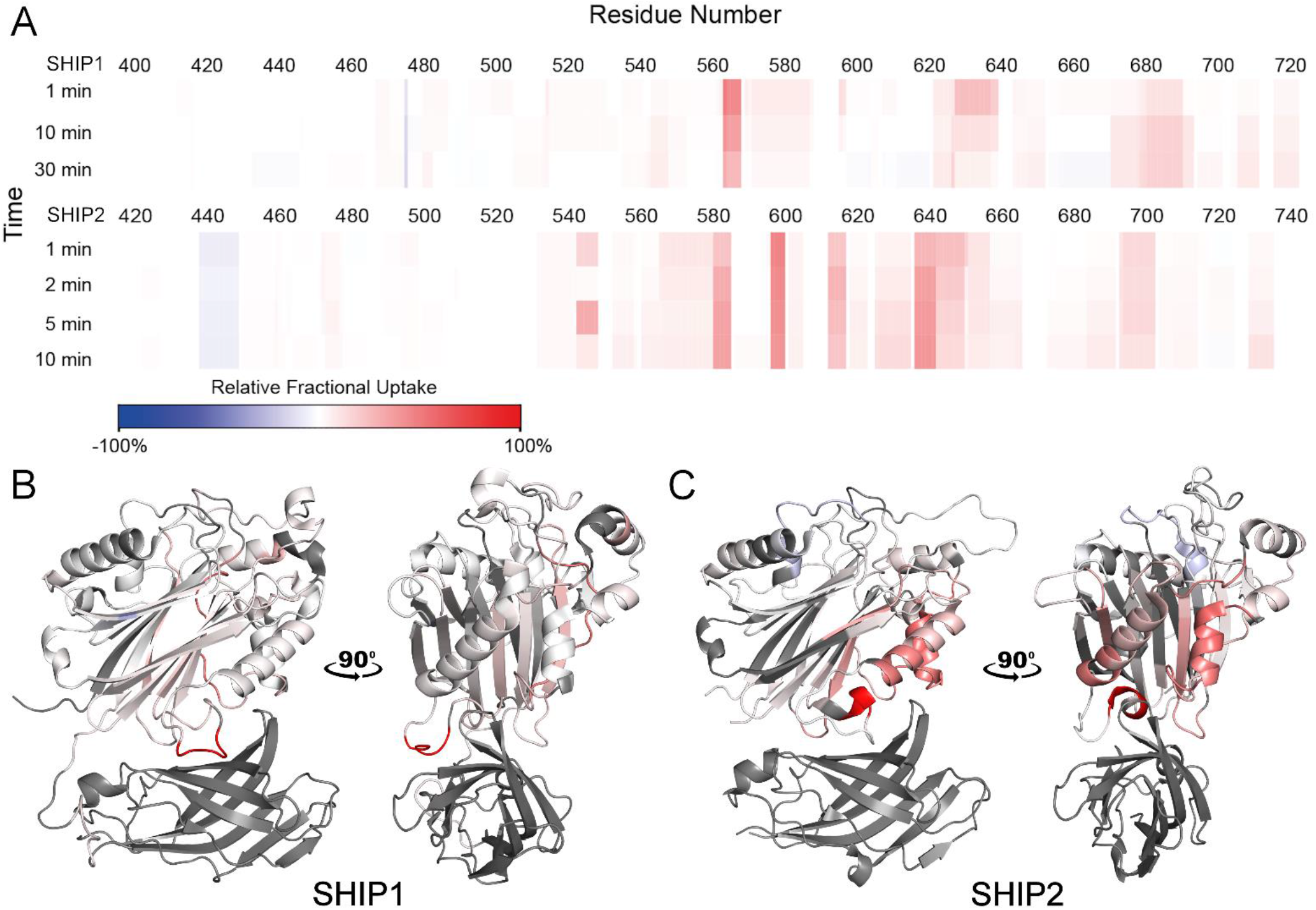
Hydrogen deuterium exchange. **(A)** Differential heat maps showing difference in deuterium uptake (DU) between SHIP1 and SHIP2 constructs with and without the C2 domain. Greater DU for the construct with the C2 domain is shown in blue while greater DU for the construct without the C2 domain is shown in red. Constructs without the C2 domain take up more deuterium and this effect is greater in SHIP2 compared to SHIP1. **(B)** The uptake for the 1 minute time point mapped on to the structure of SHIP1 (6XY7). Regions not observed and the C2 domain are shown in grey. **(C)** The uptake for the 1 minute time point mapped on to the structure of SHIP2 (5OKM chain B).

### Molecular dynamics

We conducted molecular dynamics to compare the relative flexibility of the SHIP1 and SHIP2 phosphatase domains in either the presence or absence of the C2 domain. Initially, we aimed to replicate the molecular dynamics simulations performed by Le Coq et al.^33^ on SHIP2 with and without the C2 domain starting from a loop 4-out conformation (Figure 5A,B) and complemented these with simulations on SHIP2 starting from a loop 4-in conformation (Figure 5C,D). We also performed simulations on SHIP1 starting from a loop-4 in conformation (Figure 5E,F). Simulations could not be performed on SHIP1 starting from a loop 4-out conformation as no such structure exists. These simulations were analysed in two ways: firstly, we determined root mean square fluctuations (RMSFs) for C-alphas within the phosphatase domain, examining the effect of the C2 domain on the flexibility of the phosphatase domain, and particularly loop 4. Secondly, we calculated minimum distances across the simulations between K664/R682 (SHIP1/SHIP2) in loop 4 and neighbouring acidic residues (D592 in SHIP1, D613 and D615 in SHIP2) that appear to stabilise the loop 4-out conformation, which were proposed by Le Coq et al.^33^ to be residues that are involved in the signal network between the C2 domain and the active site.

**Figure 5.**
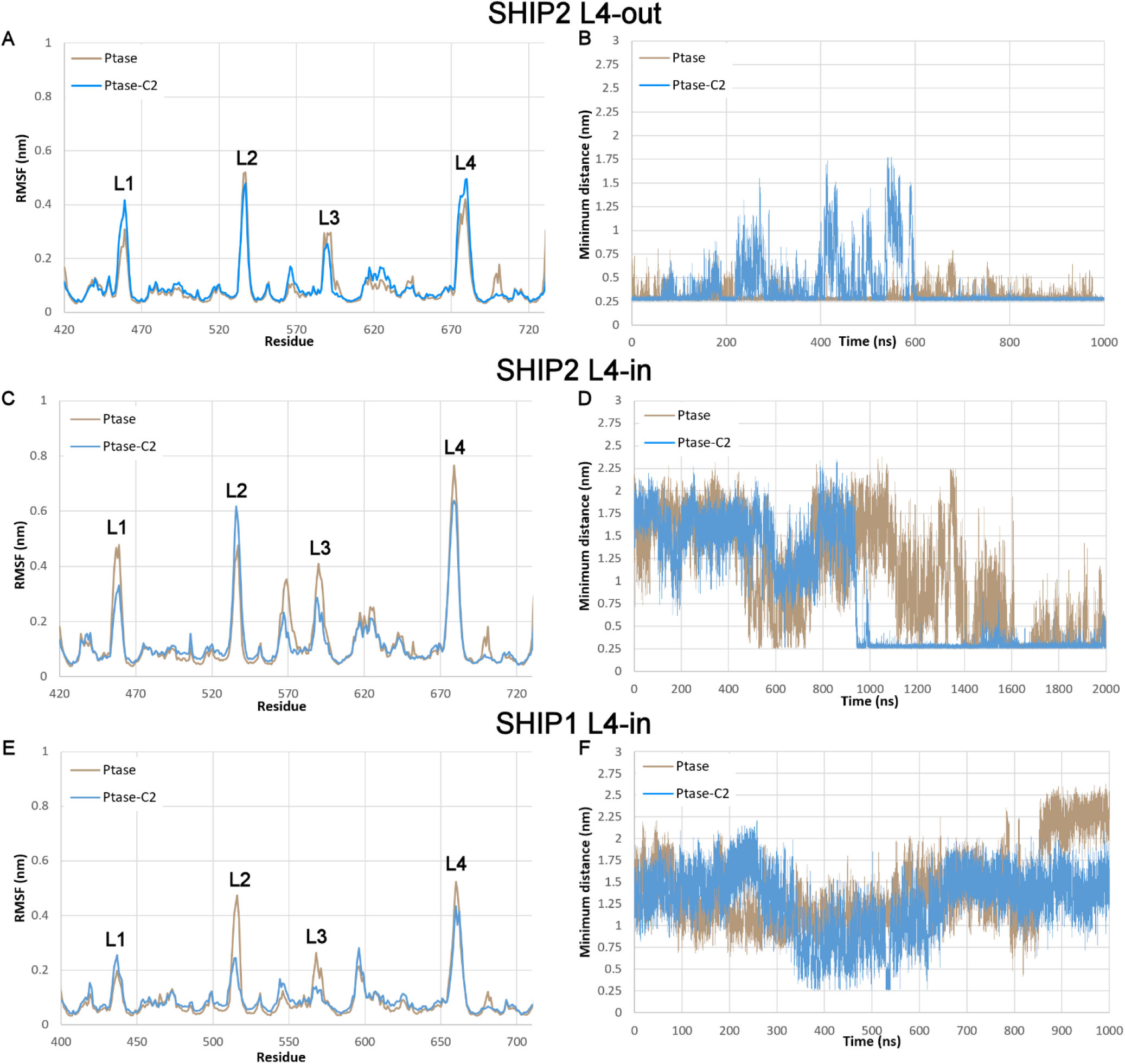
Molecular dynamics simulations. **(A)** SHIP2 L4-out. Our average (n=3) RMSFs (Root mean square fluctuations) from reproductions of the simulations performed by Le Coq et al.^33^. This figure is equivalent to figure 5A in Le Coq et al.^33^ Loops 1 to 4 are labelled. **(B)** SHIP2 L4-out. Minimum distance between R682 and D613/D615 from reproductions of the simulations performed by Le Coq et al.^33^. This figure is equivalent to figure 5D in Le Coq et al.^33^ and is a representative graph for 3 repeats. **(C)** SHIP2 L4-in. Average RMSFs from simulations using 3NR8^31^ and 3NR8 with the C2 domain from 5OKM chain B. **(D)** SHIP2 L4-in. Representative minimum distance between R682 and D613/D615 from simulations using 3NR8 and 3NR8 with the C2 domain from 5OKM chain B. **(E)** SHIP1 L4-in. Average RMSFs from simulations using the present apo structure, 6IBD. **(F)** SHIP1 L4-in. Representative minimum distance between K664 and D592/E597.

Our simulations on SHIP2 starting from a loop 4-in conformation showed the same general trends as those performed by Le Coq *et al.*, i.e., loop 4 is more flexible and the hydrogen bonds stabilising an L4-out conformation are more easily lost when the C2 domain is present, however, this effect was not as pronounced in our simulations. The simulations on SHIP2 that started from a loop 4-out conformation indicated the opposite: that loop 4 is more flexible and the hydrogen bonds stabilising an L4-out conformation are more easily lost when the C2 domain is *not* present. The simulations on SHIP1 from a loop 4-in conformation showed mixed results, the C-alpha RMSFs suggested loop 4 is more flexible without the C2 domain while the minimum distances between K664 and D592 suggested that the presence of the C2 domain does not affect the ability of hydrogen bonds to form between these residues.

### Crystallography-based Fragment screen

To provide a starting point for the design of novel SHIP1 ligands, we performed a crystallography-based fragment screen on crystals of the phosphatase and C2 domains of SHIP1 grown in the same conditions as the magnesium and phosphate-bound structure. Of the 597 fragments soaked into SHIP1 crystals, clear density was observed in PanDDA^35, 36^ event maps for 91 compounds in 108 binding events (Figure 6) spread across 12 sites. 85 of these binding events occurred at one of two sites either side of the phosphatase domain. As expected, no fragments were observed to be bound to the highly polar active site and the binding of many of the fragments was likely stabilised by crystallographic symmetry. There were, however, five fragments (5RWD, 5RWL, 5RXV, 5RY9 and 5RYC) bound near to the interface between the phosphatase and C2 domains, including two fragments (5RXV and 5RYC) covalently bound to Cys505 (Figure 6C), which is not conserved in SHIP2. This finding presents a potential avenue for selectively targeting SHIP1 and modulating the interaction between the two domains and therefore possibly altering the activity of the phosphatase domain.

**Figure 6.**
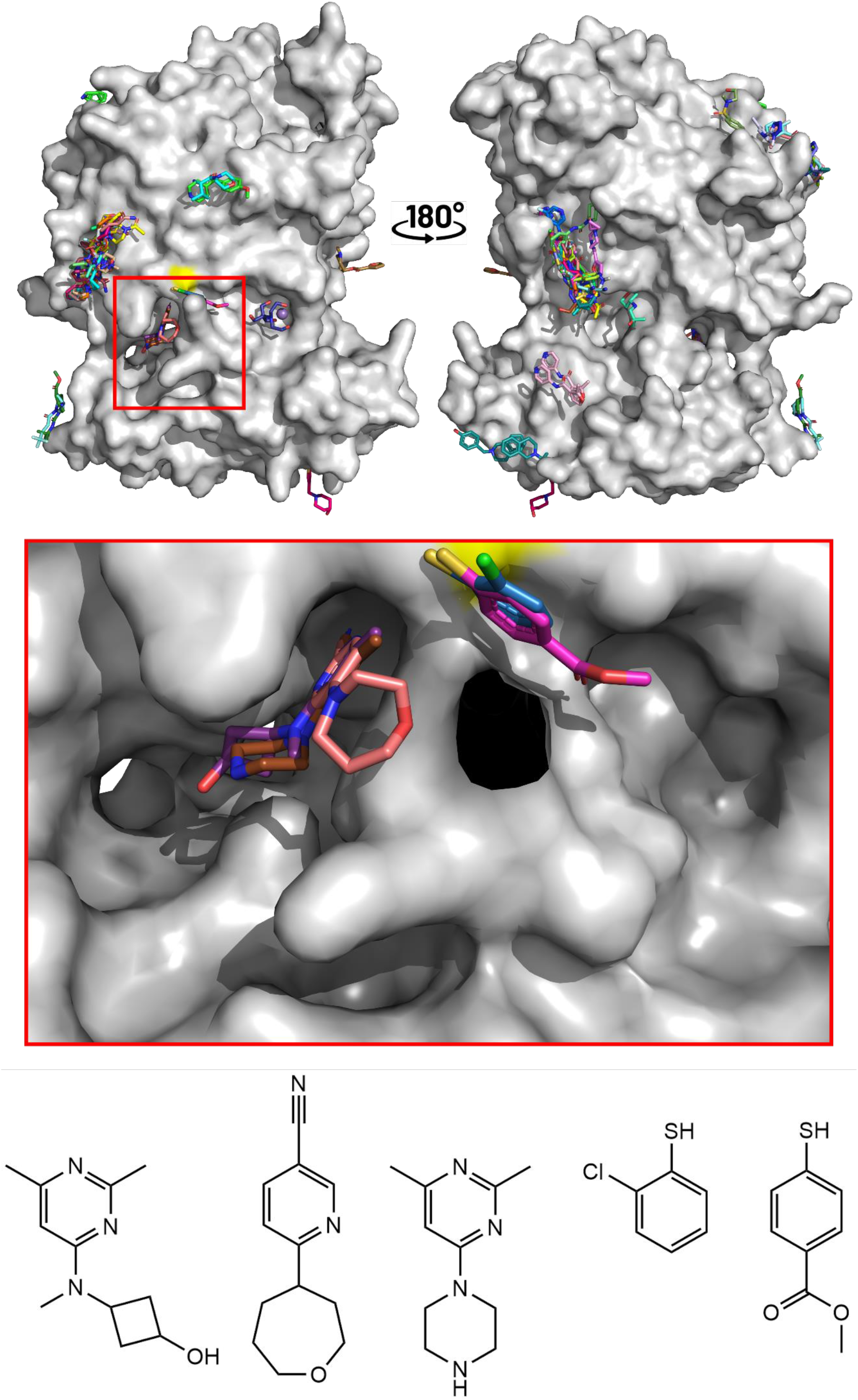
Crystallographic fragment binding. 108 binding events were observed for 91 fragments. The majority are likely to be crystallographic artefacts, but a small number were seen near to the interface between the phosphatase and C2 domains (insert), including two fragments covalently bound to C505. It may be possible to use these fragments as a starting point to disrupt the allosteric interaction between the two domains. Lys567 has been omitted from the insert for clarity. The five fragments bound near C505 are shown below, with the two covalent fragments positioned on the right.

### Mass spectrometry-based screen

As two fragments were observed to be covalently bound to SHIP1 in the crystallography-based screen, we conducted a mass spectrometry-based screen using an in-house library of 1920 covalent compounds (Figure 7A). Briefly, the protein was incubated overnight in the presence of each of the compounds and binding was assessed on an LC-ESI-TOF instrument. The results were inspected visually and the 20 strongest binders were selected for crystal soaks. Of these 20 compounds, 4 were observed to be covalently bound to Cysteine 505 in the resulting structures (Figure 7B, Table S1). For Z56948267 (8PDJ), no density was observed for the nitrile group or the chloroacetyl group, which appear to have been lost.

**Figure 7.**
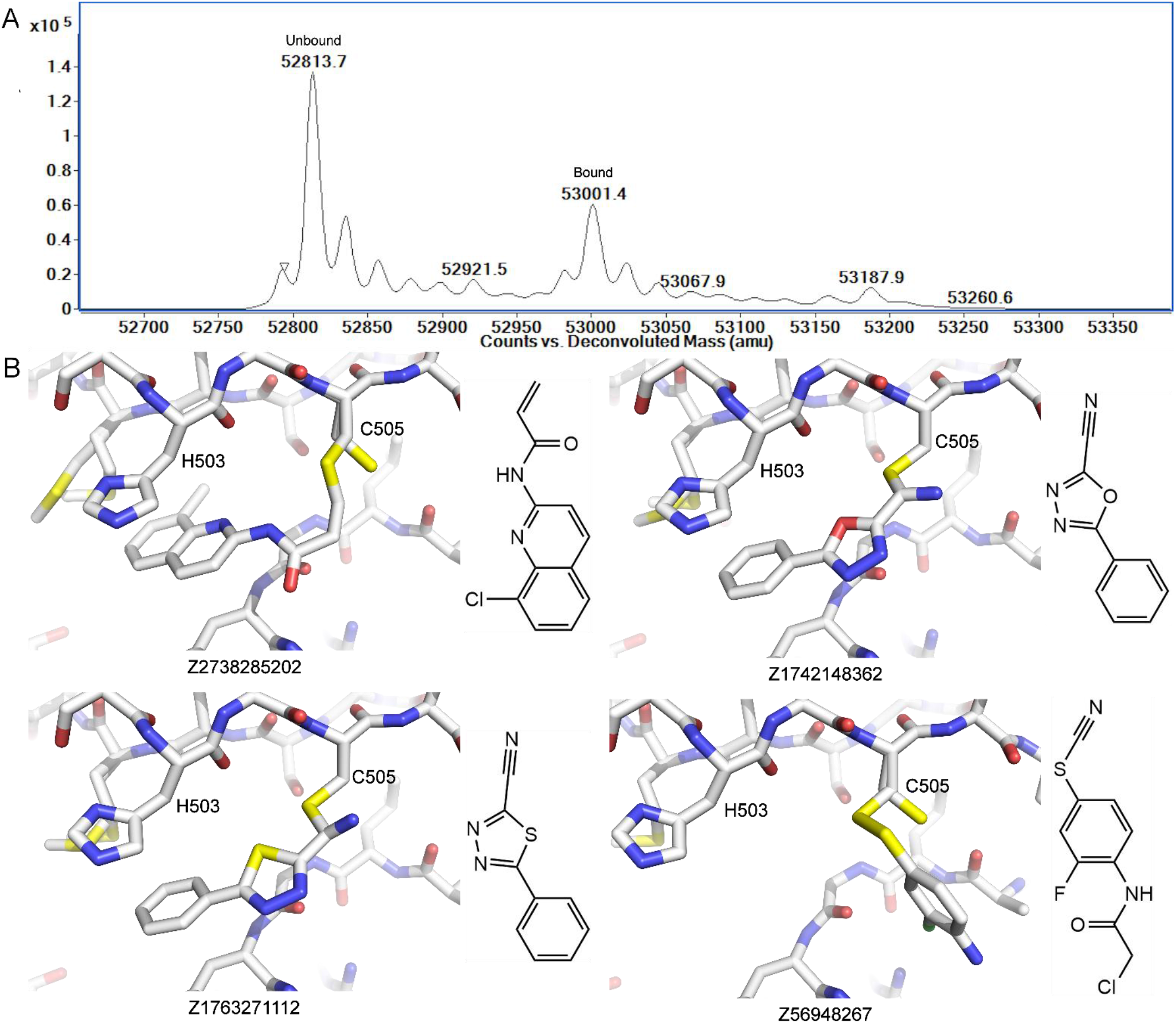
Mass spectrometry screen. **(A)** An example spectrum from the screen is shown. The unbound protein has a mass of 52813 and shows sodium adducts. The protein with a covalent ligand bound has a mass of 53001. **(B)** The four compounds that were successfully soaked into SHIP1 crystals covalently bound to C505. To the right of each image, the structure of the unbound compound is shown. In the case of Z56948267, both the nitrile group and the chloroacetyl group were lost.

## Discussion

The presented structures of SHIP1 provide a basis for comparison to SHIP2 and other inositol 5-phosphatases, highlighting potential differences that could be explored for the development of selective compounds to modulate their activity. While the overall structures of SHIP1 and SHIP2 are very similar, there are some potentially important differences. The structures of the SHIP2 phosphatase and C2 domains (5OKM, 5OKN, 5OKO, 5OKP) are in three different space groups and contain up to 8 molecules in the asymmetric unit, offering 11 “snapshots” of the structure.^33^ The two proteins appear to differ more in the C2 domain than the phosphatase domain. Respective domains of the two proteins superpose with RMSDs between 0.38 Å and 0.41 Å for the phosphatase domain, and 0.67 Å and 0.82 Å for the C2 domain. In addition, relative orientations of the phosphatase and C2 domains appear to differ somewhat between SHIP1 and SHIP2. This difference appears to result from a hinging around the linker between the two domains. When the phosphatase domains are superposed, residues in the C2 domains near to the linker superpose well, but more distant residues do not. The C-alphas of Asp809 (SHIP1, MgPO_4_-bound) and Asp829 (SHIP2, 5OKM chain G), for example, on the opposite side of the protein, are 3.2 Å apart. This hinging movement is supported by normal mode analysis with CCP4MG^37^ indicating flexibility of the C2 domain. This flexibility is further reflected in the higher B-factors observed in the C2 domains compared to the phosphatase domain in the presented structures of SHIP1 as well as those of SHIP2.

### Substrate binding and catalysis

Inositol 5-Phosphatases have been mechanistically compared to the distantly related homologues apurinic/apyrimidic (AP) endonucleases.^31, 33^ Competing hypotheses exist for the mechanism of AP endonucleases: either one metal ion^38^ or two metal ions^39^ may be required for catalysis and, if the mechanism uses a single metal ion, this ion may move to different sub-pockets of the active site during the reaction (“site A” and “site B”). Recent developments on these hypotheses are summarised by Miroshnikova et al.^40^ It remains unclear, however, which is correct and to what extent this can be extrapolated to inositol 5-phosphatases. Regardless of the number of metal ions required for the action of inositol 5-phosphatases and whether those metal ions move during the reaction, it seems that the first step of the reaction is a nucleophilic attack on the 5-phosphate by a water deprotonated by Asp586/Asp607 (SHIP1/SHIP2). The magnesium and phosphate-bound structure of OCRL (4CMN, Figure 2C) is believed to demonstrate the pre-catalytic positioning of the leaving phosphate with enough space between it and the catalytic Asp422 to accommodate the attacking water.^31^ This water, however, is not visible in the structure, likely due to the resolution of 3.13 Å. This resolution also limits the precision of the placement of the phosphate ion, the magnesium ion, and a coordinating water. Conversely, there is not enough space for this water in our magnesium and phosphate bound SHIP1 structure (Figure 2A), or the equivalent structure of SHIP2 (5OKN, Figure 2B). There is a direct interaction between Asp586/Asp607 and the phosphate, which in SHIP1 is approximately 2.3 Å away from its position in the OCRL structure. It was suggested that this represents the post-cleavage position of the 5-phosphate.^33^ Which our higher-resolution SHIP1 structure seems to confirm.

The structure of the 5-phosphatase domain of synaptojanin-1 was recently determined with PI(3,4,5)P_3_-diC8 bound to the active site.^32^ This is the first substrate-bound inositol 5-phosphatase structure. Despite a relatively low resolution of 2.73 Å, the inositol ring, phosphates, substrate glycerol moiety, and most active site residues have good density. The density for the acyl tails, magnesium ion and coordinating waters is somewhat lacking, so slightly limits the conclusions that can be drawn and the Check My Metal Server^41^ returns poor validation metrics for the magnesium ion. The relative orientation and distance between the magnesium ion and the inositol ring suggests the possibility of a hydrated cation–π interaction.^42^ The leaving phosphate is closer to the position seen in the present SHIP1 structure than in the OCRL structure, i.e., post-cleavage rather than pre-cleavage, despite the structure being substrate-bound the potential attacking water is oddly positioned. The magnesium ion, on the other hand, is closer to the position seen in the OCRL structure than in the SHIP1 structure. Further high-resolution and substrate-bound structures will be required to fully explain these differences and characterise binding and the catalytic mechanism.

There are three notable differences between inositol 5-phosphatase active sites (Figure 2). Firstly, Gly413/Gly434 in SHIP1/SHIP2 is usually an asparagine in other INPP5s, but may also be a glutamine or alanine. Secondly, Ser543/Ser564 is usually an alanine, but may also be a histidine or proline. Additionally, loop 4 in the SHIPs is replaced with the shorter P4-interacting motif (P4IM). The first two of these differences are particularly notable since, in the OCRL and synaptojanin-1 structures, the asparagine coordinates the magnesium ion. As this coordination donor is not present in SHIPs, since they contain a glycine at this position, any “pull” on the magnesium towards that specific part of the active site is lost, arguably resulting in the different binding of magnesium and phosphate seen in the SHIP structures. Instead, the phosphate is now weakly bound to the serine. The inference of weak binding is taken from the fact that this serine has clear density for 3 different conformations in our high-resolution structure, two of which place the oxygen 2.5 Å and 2.7 Å away from one of the phosphate oxygens. If it is argued that the positioning of the magnesium ion in the SHIP structures represent the post-catalytic position and the position in the OCRL and synaptojanin-1 structures represent the pre-catalytic position, how would the magnesium get into the precatalytic position in the SHIPs, as it must do for catalysis to proceed? Does the substrate force the magnesium into the precatalytic position? Trésaugues et al.^31^ observed that SHIP2 is between 25 and 80 times less catalytically active than other inositol 5-phosphatases. It was suggested that this may simply be because the assay was not optimised for SHIP2. While their assay only used the phosphatase domain of SHIP2, which both we and Le Coq et al.^33^ have shown is considerably less active without the C2 domain, we have observed similarly lower levels of activity for both SHIP1 and SHIP2, comparable to that seen by Trésaugues et al.^31^. Thus, we propose that the differences in the active site favouring the binding of magnesium, and therefore phosphate, in the “product position” in the SHIPs, as opposed to OCRL which favours the “substrate position”, may be the reason for their lower activity. There also could be a degree of coevolution in these two residues (Figure 2F), suggesting that their roles in the mechanism are linked.

The third of the differences – the 11-residue loop 4 in SHIPs versus the 7 residue P4IM in other inositol 5-phosphatases – may play an important role in substrate specificity. The P4IM is so named because it closely interacts with the phosphate bound to position 4 of the inositol ring.^31^ The longer loop 4 still interacts with the 4-phosphate but is believed to fold over the top of the substrate, allowing K664/R682 to interact with the phosphate in position 3.^33^ While the SHIPs can cleave substrates without a 3-phosphate, they seem to be considerably more active when the 3-phosphate is present. The role of loop 4 in conveying the allosteric effect from the C2 domain may explain this selectivity.

It has recently been shown that while the primary biological substrates of SHIP1 are I(1,3,4,5)P_4_ and its phosphatidyl phosphodiester, PI(3,4,5)P_3_, the protein is able to hydrolyse a range of substrates.^43^ Likewise, SHIP2 has been shown to be capable of hydrolysing several (phosphatidyl) inositol phosphates,^44^ but notably, SHIP1 was shown to hydrolyse I(2,3,4,5)P_4_, whereas SHIP2 cannot. Nelson et al.^43^ were unable to explain this observation. The active site is well conserved between the two SHIPs, except for a number of residues in loop 4 and N414/S435. The side chain of N414 seems to be well positioned to act as a hydrogen bond donor to a phosphate in position 2. It seems unlikely that any residues in loop 4 will be responsible for this substrate selectivity, so N414/S435 is the primary candidate. Regardless of the structural basis for this selectivity, this distinction may be the sole difference between the activities of SHIP1 and SHIP2, which could potentially be exploited in the development of selective inhibitors that bind to the active site.

### The Allosteric mechanism

The allosteric effect of the C2 domain on phosphatase activity was explored in SHIP2 by Le Coq et al.^33^ using a malachite green assay. It was observed that for I(1,3,4,5)P_4_, the k_cat_ of a construct possessing the phosphatase and C2 domains was approximately 50% higher than for the phosphatase domain alone, while the K_m_ was marginally higher when the C2 domain was present. They also observed that for PI(3,4,5)P_3_-DiC8, the presence of the C2 domain resulted in an 11-fold increase in k_cat_ and a doubling of K_m_. We carried out kinetic characterisation for the SHIP1 phosphatase domain in the presence and absence of the C2 domain for I(1,3,4,5)P_4_ and PI(3,4,5)P_3_-DiC8 (Figure 3A). The same characterisation was performed for SHIP2 (Figure 3B). In concordance with data shown by Le Coq et al.^33^ the C2 domain significantly enhances the catalytic activity of the phosphatase domain of SHIP2 and this effect is more pronounced for PIP_3_ than for IP_4_. In contrast, the C2 domain of SHIP1 appears to cause a moderate increase in catalytic activity for PIP_3_ but has no significant effect on activity for IP_4_.

Le Coq et al.^33^ further characterised the allosteric effect through crystallisation of SHIP2 and activity assays on a series of mutants and molecular dynamics. It was concluded that the allosteric effect of the C2 domain is transmitted through Phe855 and Glu862 in the C2 domain, which interact with Arg649 just after helix 7. The allosteric affect then travels up the side of the phosphatase domain via helices 7, 6 and 5 to a hydrogen bond network involving Asp613, Asp615 just before helix 5 and Arg682 in loop 4 (Figure 8). Loop 4 forms part of the active site, allowing the C2 domain to modulate the flexibility of loop 4 between “L4-in” and “L4-out” conformations, thereby affecting substrate binding and product release. These charge-based interactions were determined to mainly affect interactions with the polar inositol head group, while less characterised hydrophobic interactions were said to affect binding to the acyl tails.

**Figure 8.**
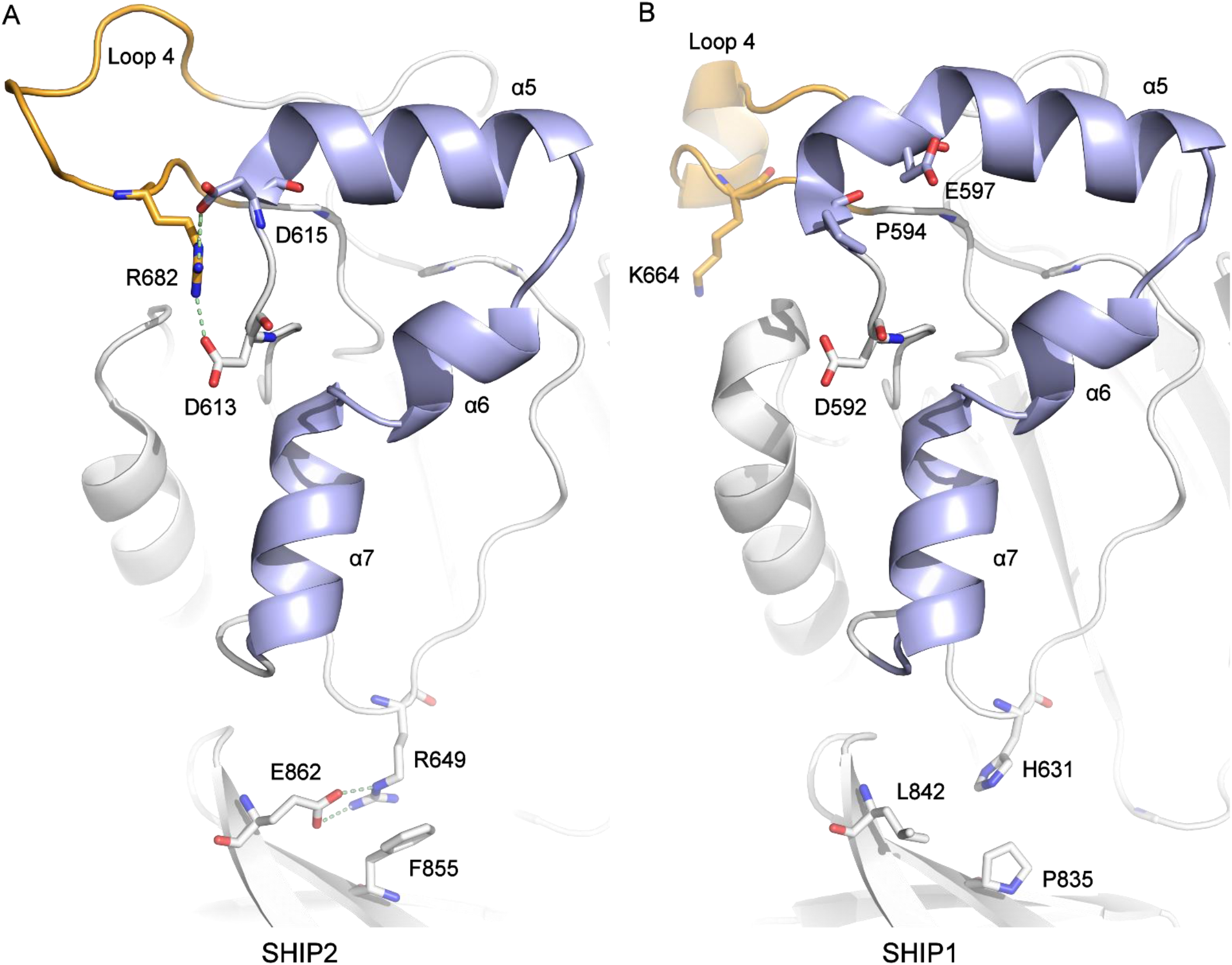
The proposed allosteric mechanism. **(A)** SHIP2 L4-out. Le Coq et al.^33^ proposed that the effect of the C2 domain is conveyed from Phe855, Glu862 to Arg649, up helices 7, 6 and 5 (light blue), through Asp613 and Asp615 to Arg682 in loop 4 (orange), which is central to substrate binding. **(B)** SHIP1 L4-in. These residues are largely unconserved in SHIP1, leading to the absence of the required interactions.

It seems logical that the allosteric mechanism would be conserved between SHIP1 and SHIP2, the phosphatase domains of which have 63% identity, while the C2 domains have 43% identity. However, the first three residues identified in the allosteric mechanism, Phe855, Glu862 and Arg649, are not conserved (Figure 8). Of the latter three residues, Asp613/Asp592 (SHIP2/1) is conserved, while Arg682 (SHIP2) is exchanged for Lys664 (SHIP1). Initially, this may seem mechanistically acceptable, but lysine is unable to form the number of hydrogen bonds hypothesised as important. Asp615 (SHIP2) is not conserved (Pro594 in SHIP1), but it is possible that its role is fulfilled by the nearby Glu597 (SHIP1). As our structures are in a loop4-in conformation, they do not show hydrogen bonds between Lys664 and either Asp592 or Glu597. To shed further light on the allosteric mechanism and possible differences between SHIP1 and SHIP2, we replicated the molecular dynamics performed by Le Coq et al on SHIP2 starting from a loop 4-out conformation and performed the same simulations on SHIP2 starting from a loop 4-in conformation, and on SHIP1 from a loop-4 in conformation. No structures of SHIP1 in a loop 4-out conformation have been published yet, so it was not possible to perform these simulations. We also performed hydrogen-deuterium-exchange mass spectrometry to investigate changes in the flexibility of the phosphatase domain caused by the absence of the C2 domain.

Our replication of the MD performed by Le Coq *et al.* showed the same general trend, ie. loop 4 is more flexible and the hydrogen bonds stabilising an L4-out conformation are more easily lost when the C2 domain is present. This effect was not as pronounced in our simulations, however. Starting the SHIP2 simulations from a loop 4-in conformation indicated the opposite: that loop 4 is more flexible and the hydrogen bonds stabilising an L4-out conformation are more easily lost when the C2 domain is *not* present. While simulations on SHIP1 from L4-in indicated that the flexibility of loop 4 largely is unaffected by the presence of the C2 domain.

In the SHIP1 molecular dynamics simulations, regardless of the presence of the C2 domain or a hydrogen bond, Lys664 was always closer to Asp592, than Glu597, which is not the case for the respective SHIP2 residues, suggesting that Glu597 in SHIP1 does not fulfil a role equivalent to that of Asp615 in SHIP2 and that the exchange of Arg682 for Lys664 and Asp615 for Pro594 results in a partial loss of the hydrogen bond network seen around loop 4 when SHIP1 is compared to SHIP2.

Our HDX-MS provides further insight into the allosteric mechanism proposed by Le Coq *et al.*, who said that the C2 domain is able to modulate the flexibility of loop 4 via helices 5, 6 and 7. The HDX-MS clearly demonstrated that in SHIP2, when the C2 domain is not present, loop 3, strand 9 and helices 5, 6 and 7 had increased deuterium uptake, indicating increased solvent accessibility and/or flexibility. This effect was also seen in SHIP1 but to a lesser extent. Loop 4 showed maximal deuterium uptake at the first time point for both SHIP1 and SHIP2. This seems to indicate that the effect of the C2 domain is conveyed to the opposite side of the phosphatase domain by helices 5, 6 and 7 but also by strand 9. Residues identified by Le Coq *et al.* may be involved in this mechanism and their lack of conservation may explain why the effect appears to be weaker in SHIP1. It seems that the C2 domain causes a significant degree of stabilisation and decreased flexibility for one side of the phosphatase domain. Our molecular dynamics simulations do seem to reflect this to some extent. It does seem that, when starting from a loop 4-in conformation, at least, the phosphatase domain is marginally more flexible when the C2 domain is not present (Figure 5C,E). Although this was not seen for the SHIP2 L4-out simulations (Figure 5A). The lack of effect either way seen in SHIP1 may reflect the decreased effect of the C2 domain.

The results from MD, combined with the results from HDX-MS and the lack of conservation of important residues between SHIP1 and SHIP2 seem to indicate that while the allosteric mechanism hypothesised by Le Coq *et al.* is likely to hold some validity. Specifically, there appears to be some altered flexibility of helices 5, 6 and 7, possibly influenced by the ability of the C2 domain to modulate the flexibility of one side of the phosphatase domain, potentially through a hinging motion around the linker connecting the two domains. This modulation may impact the flexibility of loop 4 although our HDX-MS experiments did not reveal significant changes and the observed changes in MD simulations were minimal. Notably, much of the hydrogen bond network around loop 4 in SHIP2 that was proposed to be involved in the mechanism is not conserved in SHIP1. The apparent weaker effect of the C2 domain on the phosphatase domain of SHIP1 may mean that modulation of phosphatase activity by targeting the C2 domain is perhaps a more viable strategy for SHIP2 than for SHIP1. However, it is important to not entirely disregard the potential of targeting the C2 domain of SHIP1 as it does seem to still have some effect.

Further to the work presented here, it has been demonstrated recently that the PH domain that precedes the phosphatase domain in both SHIP1 and SHIP2 is also able to modulate the activity of the phosphatase domain, in SHIP2, at least.^45^ It has also been shown that the compound ZPR-MN100 is able to activate SHIP1, probably through binding to a pocket at the interface between the phosphatase and C2 domains, likely modulating the interaction between the two domains with Lys681 (mouse residue numbering, Lys677 in human) being essential to binding.^34^ Further investigation into the mechanisms involved in both would potentially unlock the opportunities target the two domains either side of the phosphatase domain as strategies for developing small molecule modulators.

### Fragment-bound structures

To provide starting points for the development of SHIP1 modulating compounds, we have presented 95 structures of SHIP1 with small molecules bound with 91 derived from crystallography-based fragment screening and 4 from a mass spectrometry screen. One of two sites at which the majority of binding events occurred in the crystallographic screen is in close proximity to the Lys677 pocket, to which it has been suggested ZPR-MN100 binds.^34^ These compounds may provide potential ways in which this compound could be elaborated. We also observed five molecules from the crystallographic screen bound to a pocket near to the interface between the phosphatase and C2 domains, of which 2 were covalently bound to Cys505, a residue not conserved in SHIP2. The 4 structures that contain compounds identified in the MS screen show the molecules covalently bound to the same cysteine residue. It is assumed that other hits from the MS screen that we were unable to determine structures for also bound to this cysteine. These structures present promising starting points for the development of potent and selective molecules able to bind to SHIP1. Many of the compounds identified in this study have been assayed for their ability to modulate SHIP1 activity (data not shown) but so far, all show minimal effects and should only be taken as starting points. Given their proximity to the interface between the two domains, compounds bound at this site have the potential to modulate the activity of the phosphatase domain. Interestingly, helix 4, which showed increased flexibility in the constructs without the C2 domain in HDX-MS analysis is near this cysteine, providing a mechanism through which compounds bound to Cys505 may modulate phosphatase domain flexibility and therefore, activity.

We have presented here high-resolution structures of the phosphatase and C2 domains of SHIP1 and assessed the role of the C2 domain in SHIP1 and SHIP2 using activity assays, hydrogen-deuterium-exchange mass spectrometry and molecular dynamics. While the C2 domain of both proteins can modulate the activity of the phosphatase domain through a seemingly conserved mechanism, the effect is considerably more pronounced in SHIP2 than in SHIP1, which we put down to the lack of conservation of specific residues identified as important to allosteric communication. We have also presented a crystallographic fragment screen on SHIP1 and followed this up with a mass spectrometry screen of covalent compounds, from which we were also able to determine a further four structures with covalent compounds bound. These structures will serve as starting points for the development of novel and selective compounds able to modulate the activity of SHIP1 by targeting the allosteric mechanism discussed herein.

## Methods

### Cloning

LIC primers were designed corresponding to residues Glu396 to Gln856 of human SHIP1 (SHIP1-Ptase-C2) and the pGTVL2 (used for 6IBD) and pFB-HGT-LIC vectors (used for all other work). The PCR product was ligated into the two vectors with the former transformed into Rosetta cells (BL21(DE3)-R3-pRARE2) and the latter into DH10Bac for use with the Bac-to-Bac system. SHIP2 constructs coding for Glu420 to Ser743 (SHIP2-Ptase) and Glu420 to Arg878 (SHIP2-Ptase-C2) were similar cloned into pFB-HGT-LIC. Successful DH10Bac transformants were identified through blue/white screening and the bacmid was purified from 2 ml overnight cultures supplemented with appropriate antibiotics. The SHIP1 construct coding for Asn392 to Gln729 in pS97splitRBP (SHIP1-Ptase, SHIP1ΔC2) was received from William Kerr and Sandra Fernandes and transformed into Rosetta cells.^46^

### Protein expression

#### Bacterial expression

One 50 ml and two 10 ml overnight LB cultures inoculated from glycerol stocks were grown in a 250 ml conical flask and 50 ml falcon tubes respectively, each with 50 μg/ml kanamycin and 34 μg/ml chloramphenicol at 37 °C with shaking at 250 rpm. The following morning, the cultures were combined and 10 ml was used to inoculate each of six 1L TB cultures with 50 μg/ml kanamycin. They were grown at 37 °C with shaking at 180 rpm. Once the cultures had reached an OD_600_ between 1.5 and 2.0, the temperature was reduced to 18 °C and, 30 minutes later, expression was induced by addition of 300 μM IPTG. Cultures were grown overnight and harvested by centrifugation at 6,000g the following morning. Pellets were scraped into falcon tubes and flash frozen in liquid nitrogen.

#### Insect cell expression

It was observed that some SHIP1 and SHIP2 constructs produce a significantly greater yield through insect cell expression. The constructs were expressed in Sf9 cells grown in Sf-900 II SFM. Each litre of cells at a density of 0.2×10^6^/ml was infected with 3 ml of P2 viruses and allowed to grow for approximately 68 hours at 27 °C. Cells were harvested by centrifugation at 1500g, flash frozen in liquid nitrogen and stored at −80 °C.

#### Protein purification

For bacterially expressed protein, 6 L of expression pellets were thawed by immersing the tubes in water and were resuspended in lysis buffer (50 mM HEPES pH 7.5, 500 mM NaCl, 20 mM imidazole, 5% glycerol, 1 mM TCEP) to a total volume of 240 ml per 3 L of expression pellet. For protein expressed in Sf9 cells, 2L Expression pellets handled in the same way were resuspended in lysis buffer to 240 ml supplemented with EDTA-free protease inhibitors. The cells were lysed by sonication on ice with 5 seconds on, 10 seconds off for a total of 20 minutes for *E. coli* and 15 minutes for Sf9. The lysate was mixed halfway through sonication. Once lysis was complete, the lysate was cleared by centrifugation at 75,000g for 20 minutes.

After prewashing 3 ml of nickel beads in lysis buffer, they were added to the supernatant and split into several 50 ml falcon tubes. The tubes were then rotated in a cold room for 1 hour. Subsequently, the beads were pelleted by centrifugation at 800g for 5 minutes and washed twice with lysis buffer before being pelleted again.

The beads were resuspended in 20 ml lysis buffer and loaded on to a gravity column. The resin was washed with 10 ml wash buffer (50 mM HEPES pH 7.5, 500 mM NaCl, 40 mM imidazole, 5% glycerol, 1 mM TCEP) before the protein was eluted with three 10 ml washes with elution buffer (50 mM HEPES pH 7.5, 500 mM NaCl, 300 mM imidazole, 5% glycerol, 1 mM TCEP).

TEV protease was added to the eluate at a concentration (mg/ml) of 5:1 target protein:TEV. The mixture was dialysed overnight into dialysis buffer (50 mM HEPES pH 7.5, 500 mM NaCl, 5% glycerol, 1 mM TCEP) at 4 °C. The TEV protease was removed by passing the sample back down the nickel column, equilibrated with lysis buffer, followed by a wash with 10 ml lysis buffer. The flow through and wash fractions were combined and concentrated to a volume of 1 ml. This was passed down a Superdex 200 16/60 column in gel filtration buffer (50 mM HEPES pH 7.5, 250 mM NaCl, 5% glycerol, 1 mM TCEP). Selected fractions were combined and concentrated to around 6 mg/ml.

### Structure determination

#### Crystallisation

Crystallisation conditions for SHIP1-Ptase-C2 were screened at 20 °C by sitting drop vapour diffusion using 150 nl drops with ratios of 2:1, 1:1 and 1:2. Screens were set up around identified conditions and the best crystals took around 1 to 2 weeks to grow in 0.1M bis-tris pH 7.0, 14% PEG2KMME, 12% PEG3350. The crystals were cryoprotected by addition of 1 μl reservoir to the drop. The apo structure was determined from crystals grown in these conditions. These crystals were also used for seeding. Crystals from 6 drops were resuspended in a total of 30 μl reservoir solution with a seed bead. The seed stock was vortexed for 10 seconds three times with 30 seconds on ice between each. This was then diluted 100-fold in reservoir solution and conditions were rescreened with 20 nl seeds added to 150 nl drops. Crystals were observed in Molecular Dimensions Morpheus C1 (30 mM sodium phosphate, ammonium sulphate, sodium nitrate, 100 mM MES/imidazole pH 6.5, 10% PEG 20,000, 20 % PEG 500 MME) after around a week. For the magnesium and phosphate bound structure, these crystals were cryoprotected by addition to the drop of 1 μl reservoir solution supplemented with 2mM MgCl_2_.

#### Data Collection and processing

Data were collected at Diamond Light Source beamline I03. Apo data were integrated using Dials^47^ and scaled with Aimless^48^ to a resolution of 1.48 Å, while the data for the magnesium and phosphate bound structure were scaled with Staraniso^49^ and Aimless. An elliptical high-resolution cut-off of between 1.34 Å and 1.09 Å was applied. Molecular replacement was performed using Phaser^50^ with a model generated by Swiss-Model^51^ based on chain B of the structure of SHIP2 (5OKM) and refined with successive rounds of Coot^52^ and Refmac5.^53^ The apo structure was refined to an R_work_ of 0.174 and an R_free_ of 0.200 while the magnesium and phosphate bound structure was refined to an R_work_ of 0.136 and an R_free_ of 0.157. The final models were verified with MolProbity.^54^ The magnesium ion was verified with Check My Metal^41^ and the structures were deposited in the PDB under accession codes 6IBD and 6XY7.

#### Activity Assays

The phosphatase activity of SHIP1 and SHIP2 was determined by measuring inorganic phosphate production using Phosphate Sensor (ThermoFisher Scientific, PV4407). Soluble inositol 1,3,4,5-tetraphosphate (I(1,3,4,5)P_4_) and phosphatidylinositol 3,4,5-trisphosphate-DiC8 (PI(3,4,5)P_3_-DiC8) were used as substrates. Briefly, 10 μl enzyme (SHIP1 Ptase (50 nM), SHIP1 Ptase-C2 (50 nM), SHIP2 Ptase (50 nM) or SHIP2 Ptase-C2 (15 nM) (final assay concentration)) in assay buffer (20 mM HEPES pH 7.5, 5 mM MgCl_2_, 150 mM NaCl, 0.01% Triton X-100 and 0.1% BSA) was added to clear-bottomed black 384-well plates (Greiner #781906). Phosphate Sensor, 20 μl at 1.5 μM (final concentration of 0.75 μM), was added and the plate centrifuged at 1000 rpm for 1 minute and left to incubate at RT for 30 min. Substrates, 10 μl at 4X the final assay concentration, were added to start the reaction via the internal liquid handling function of the FLIPR® Tetra (molecular devices) and the increase in fluorescence produced via phosphate binding to the Phosphate Sensor was monitored over time (λ_ex_ = 390-420 nm and λ_em_ = 475-535 nm). A dose response of the substrate was performed with a top final assay concentration of 100 μM. A phosphate standard curve was used to interpolate the concentration of phosphate produced from raw RFU values. Progress curves were normalised to the substrate alone at each concentration to account for phosphate contamination of the substrate and the initial rate of phosphate production was plotted vs substrate concentration and fitted to the Michaelis-Menten equation to obtain K_m_ and k_cat_. All data analysis was carried out in GraphPad Prism.

#### Hydrogen-deuterium Exchange

HDX-MS experiments were performed using a Synapt G2Si HDMS coupled to an Acquity UPLC system with HDX-MS and automation (Waters Corporation, UK). Labelling was performed using a continuous labelling workflow at 20 °C. Each protein was diluted to 10 μM in equilibration buffer (PBS, pH 7.4). Labelling was initiated by diluting 5 μl of each protein to 50 μl total volume by addition of labelling buffer (PBS prepared in D_2_O, pD 7.4) giving a 90% D_2_O environment. Labelling took place for various time points. The labelling reaction was quenched by a 1:1 addition of ice-cold quench buffer (50 mM KH_2_PO_4_, 1 M GdmCl, pH 2). Proteins were digested online with a Waters Enzymate BEH pepsin column at 20 °C. Peptides were trapped on a Waters BEH C18 VanGuard pre-column for 3 min at a flow rate of 200 μl/min in buffer A (0.1% formic acid ∼ pH 2.5) before being applied to a Waters BEH C-18 analytical column. Peptides were eluted with a linear gradient of buffer B (0.1% formic acid in acetonitrile ∼ pH 2.5) at a flow rate of 40 μl/min. All trapping and separation were performed at ∼0 °C to reduce back exchange.

MS data were acquired using MS^E^ workflow. All time points across this experiment were obtained in triplicate. MS was calibrated separately using NaI and the MS data were obtained with lock mass correction using LeuEnk. Peptides were assigned with the ProteinLynx Global Server (PLGS, Waters Corporation, UK) software and the isotope uptake of respective peptides was determined using DynamX v3.0.

#### Molecular dynamics

Simulations were performed using Gromacs 2018.3^55, 56^ with the AMBER99SB*-ILDN force field.^57^ SHIP1 loop 4-in simulations used the apo structure presented here, SHIP2 loop 4-out simulations used chain B of 5OKM, SHIP2 loop 4-in simulations used the phosphatase domain of 3NR8 joined to the C2 domain from 5OKM chain B. Each of these simulations was performed with and without the C2 domain. The linker between the phosphatase and C2 domains was modelled with ModLoop.^58^ Simulations used parameters based on those in the Gromacs lysozyme tutorial ^59^. The structures were prepared for full MD runs by energy minimisation followed by 100 ps temperature, pressure and density equilibration steps. MD runs were initially performed for 1 µs. As the SHIP2 loop 4-in simulations showed hydrogen bonds forming after approximately 950 ns (Figure 5D), these simulations were extended to 2 µs.

Figures 5A, 5C and 5E were calculated from the average RMSFs across the three repeats of each simulation. Averaging distances across simulations does not make sense so the data used for Figures 5B, 5D and 5F are from a single representative simulation. Those for B were selected to most strongly agree with the conclusions reached by Le Coq et al.^33^, although the other two also agreed. The phosphatase domain data used in 5D are similar to the data from the other two repeats with hydrogen bonds able to form briefly. The hydrogen bonds only formed very briefly in the other Ptase-C2 simulations, but the data shown demonstrate this formation well. The data were similar for all six SHIP1 simulations.

#### Crystallography-based fragment screening

Fragment screening was performed using the XChem pipeline at Diamond Light Source. Several hundred crystals were produced using seeding and Morpheus C1, as described above without the addition of MgCl_2_. Several crystals grown in these conditions were screened to confirm that the diffraction was routinely of a high quality and to a high resolution. Initial screening of DMSO soaked crystals showed a deterioration in diffraction quality from around 20-25% DMSO. Fragments from the DSI-posed library^60^ were therefore soaked into crystals to a final DMSO concentration of 15% (a fragment concentration of 75 mM) using a Labcyte Echo^61^ and allowed to incubate for between 2 and 5 hours. Crystals were fished from drops with the aid of a Shifter^62^, allowing approximately 200 crystals to be fished per hour. Data were collected automatically on beamline I04-1 using X-ray centring and a rapid data collection protocol (0.48° oscillations, 0.04 s exposures, 375 images). The resulting datasets were processed using XChem Explorer and PanDDA.^35, 36^ Fragment bound structures can be viewed on Fragalysis (http://fragalysis.diamond.ac.uk) and have been deposited in the PDB using accession codes 5RW2 to 5RYL and group deposition G_1002180.

### Mass spectrometry-based screen

#### Sample preparation

An in-house library of 1920 covalent small molecule fragments was dispensed in 10 nl aliquots between 7 × 384 well microplates. 12.5 µL of SHIP1-Ptase-C2 in 50 mM HEPES pH 7.5, 250 mM NaCl, 5% glycerol and 1 mM TCEP was added to each well at a concentration of 85 μg/mL to give a final ligand concentration of approximately 0.2 mM. The reaction was left overnight at room temperature and terminated by adding 50 µL of 0.2 % formic acid.

#### LC-MS screening

Reversed-phase HPLC was performed in-line prior to mass spectrometry. A 50 μl aliquot with SHIP1-Ptase-C2 with covalent library compounds was injected on to a 2.1 mm × 12.5 mm Zorbax 5 μM 300SB-C3 guard column housed in a column oven set at 40 °C. The solvent system used consisted of solvent A: 0.1% formic acid and solvent B: 0.1% formic acid in methanol. Chromatography was performed as follows: initial conditions were 90 % A and 10 % B and a flow rate of 1.0 ml/min. After 15 seconds at 10 % B, a linear gradient to 80 % B was applied over 45 seconds and then to 95% B over 3 seconds. Elution then proceeded isocratically at 95 % B for 1 minute 12 seconds followed by equilibration at initial conditions for a further 45 seconds. Protein and covalent inhibitor denaturing intact mass was determined using an MSD-ToF electrospray ionisation orthogonal time-of-flight mass spectrometer (Agilent Technologies Inc. – Palo Alto, CA, USA). The instrument was configured with the standard ESI source and operated in positive ion mode. The ion source was operated with the capillary voltage at 4000 V, nebulizer pressure at 60 psi, drying gas at 350°C and drying gas flow rate at 24 L/min. The instrument ion optic voltages were as follows: fragmentor 250 V, skimmer 60 V and octopole RF 250 V. This LC-MS method was applied to each well of the 7 microplates in 384 well format. Data analysis was performed using Agilent MassHunter Qualitative Analysis.

#### Crystallographic soaks

The top 20 hits from the MS screen were selected and soaks were performed by adding the compounds in DMSO to Morpheus C1 to a final concentration of 5 mM, 20% DMSO. 1 µl of this was added to drops containing crystals and incubated at 20 °C for one day before harvesting. Data were collected and processed as previously stated.

## Acknowledgements

The authors would like to thank Diamond Light Source for beamtime (proposals lb22717, mx15433 and mx19301), and the staff of beamlines I03, I04, I04-1 and I24 for assistance with crystal testing and data collection. We thank the Alzheimer’s Research UK Oxford Drug Discovery Institute for funding (ARUK2018DDI-OX). We also thank William Kerr and Sandra Fernandes for the donation of the SHIP1-Ptase construct.

## Author Contributions

Conceptualisation: WJB, OG. Data Curation: WJB, ECK, RC, LPB. Formal Analysis: WJB, ECK, TM, LAS, EJM, OG. Funding acquisition: PEB, OG. Investigation: WJB, ECK, TM, LAS, OG. Methodology: WJB, ECK, EJM, OG. Project administration: WJB, OG. Resources: RC, VLK, LPB, PEB, OG. Supervision: WJB, RC, VLK, LPB, PEB, EJM, OG. Validation: WJB, ECK, TM, LAS, EJM, OG. Visualisation: WJB, LAS, EJM. Writing – original draft: WJB. Writing – review & editing: All authors

---

**Table S1.**
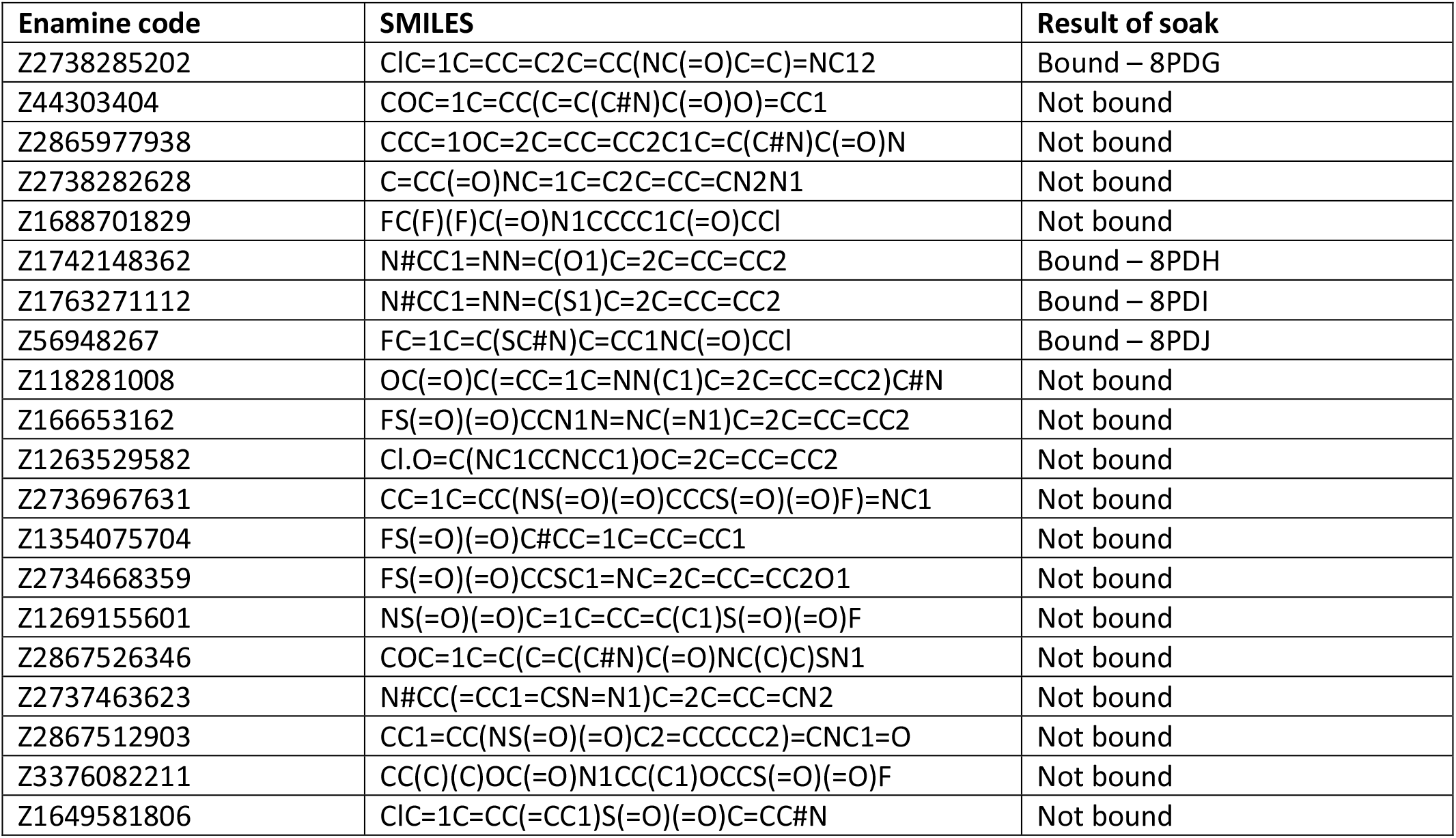
The 20 best MS screen hits.

**Figure S1.**
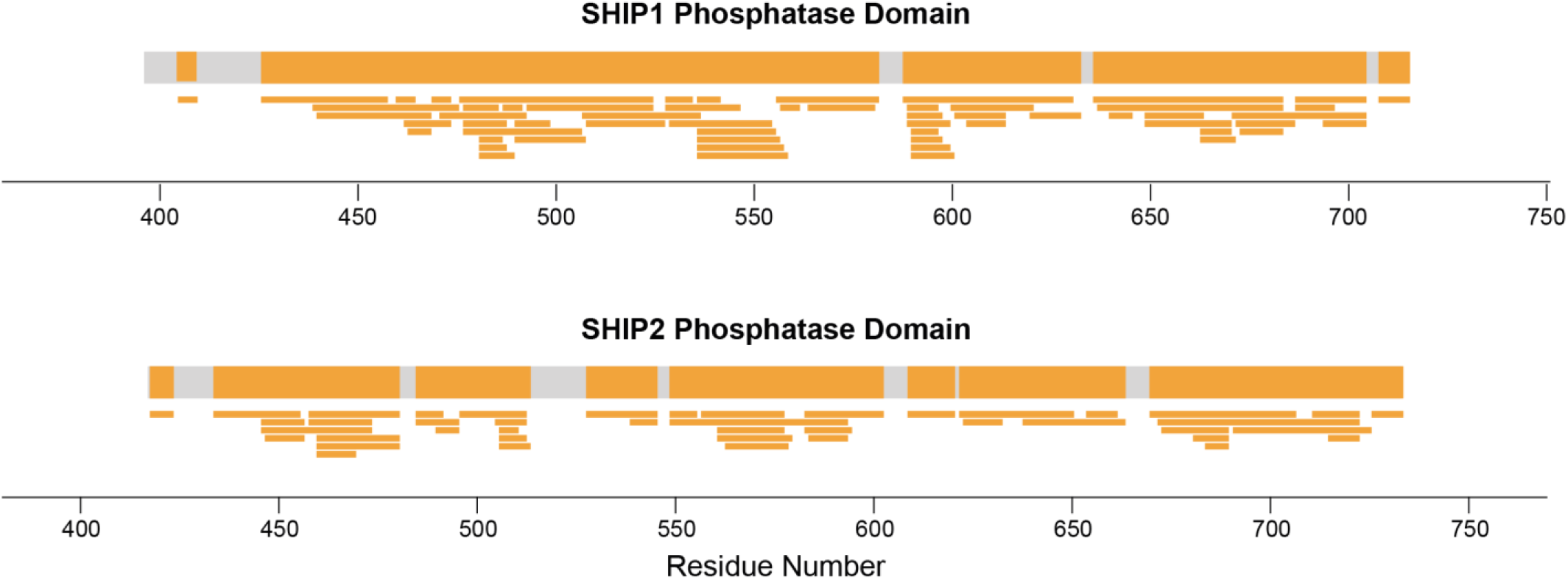
HDX-MS coverage map. Regions of the phosphatase domains of SHIP1 and SHIP2 observed in HDX-MS are shown in orange with regions not observed shown in grey. Individual detected peptides are shown below.

**Figure S2.**
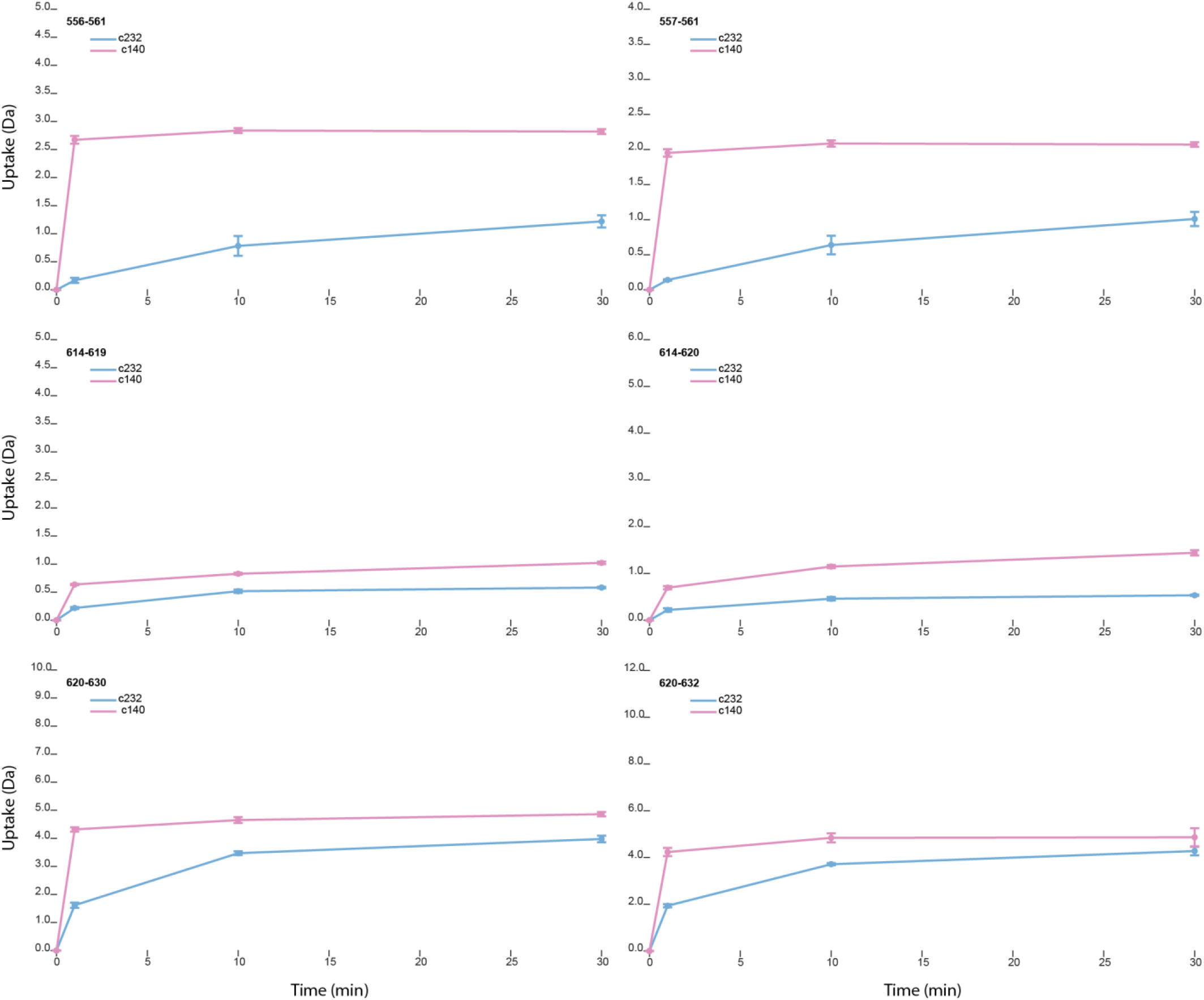
SHIP1 uptake plots. Examples of peptides detected that showed significantly greater uptake in the construct without the C2 domain than with.

**Figure S3.**
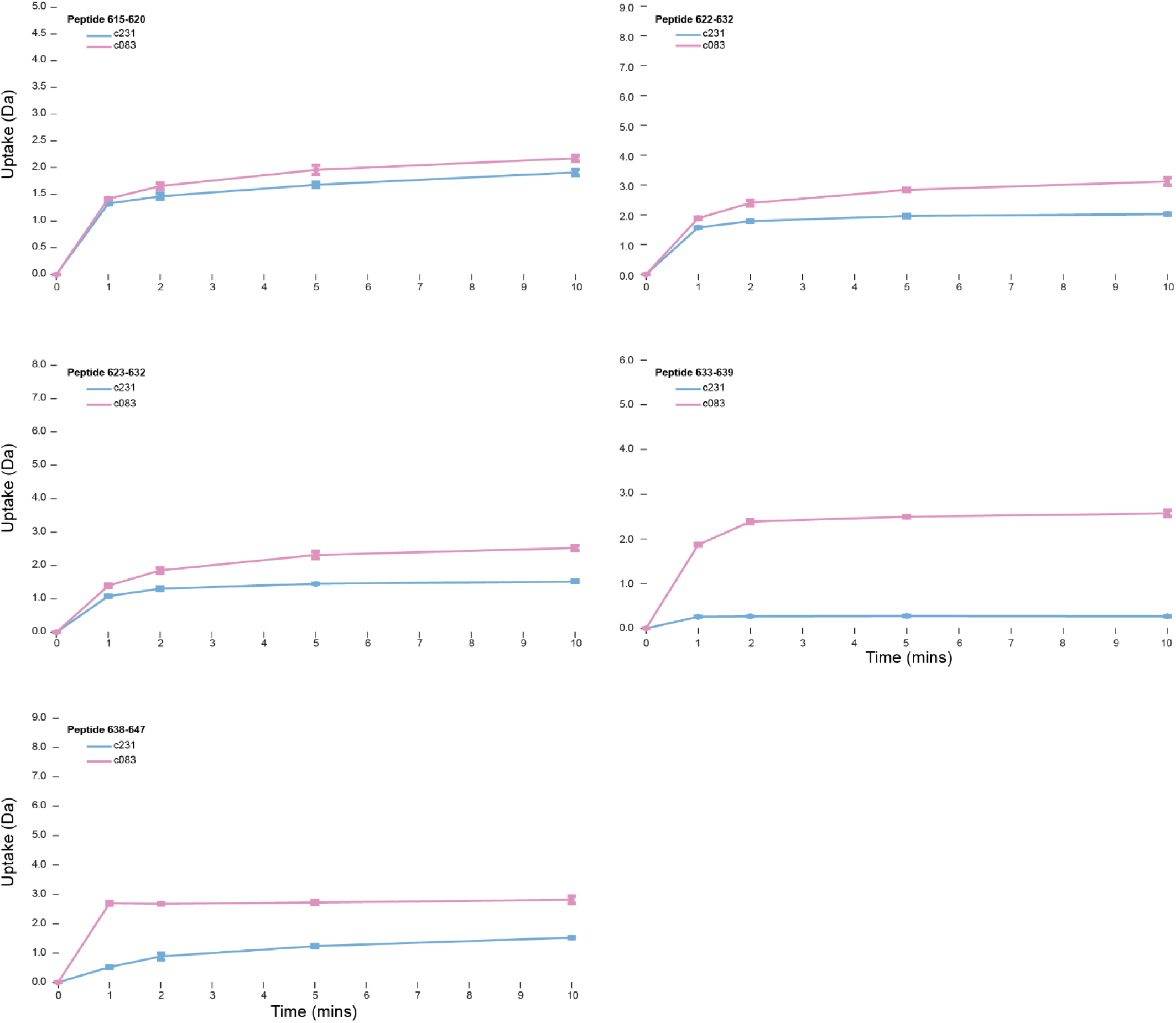
SHIP2 uptake plots. Examples of peptides detected that showed significantly greater uptake in the construct without the C2 domain than with. All plots shown are for peptides in the region of helices 5, 6 and 7.

